# A Real Time PCR Assay for Quantification of Parasite Burden in Murine Models of Leishmaniasis

**DOI:** 10.1101/312330

**Authors:** Alejandro L. Antonia, Liuyang Wang, Dennis C. Ko

**Affiliations:** Department of Molecular Genetics and Microbiology, School of Medicine, Duke University, Durham, NC 27710; Division of Infectious Diseases, Department of Medicine, School of Medicine, Duke University, Durham, NC 27710

## Abstract

Eukaryotic parasites in the genus *Leishmania* place approximately 350 million people per year at risk of disease. In addition to their global health significance, *Leishmania* spp. have served as an important model for delineating basic concepts in immunology such as T-helper cell polarization. There have been many qPCR based assays reported for measuring parasite burden in humans and animals. However, these are largely optimized for use in clinical diagnosis and not specifically for animal models. This has led several of these assays to have suboptimal characteristics for use in animal models. For example, multi-copy number genes have been frequently used to increase sensitivity, but are subject to greater plasticity within the genome and thus may confound effects of experimental manipulations in animal models. In this study, we develop a sybr-green based quantitative touchdown PCR assay for a highly conserved and single copy, putative RNA binding protein, DRBD3. With primers nearly perfectly conserved across all *Leishmania* spp., this assay rivals the sensitivity of previously reported qPCR based methods of parasite quantitation and successfully detected *L. major* from mouse infection. Use of this protocol in the future will lead to improved accuracy in animal based models and help to tease apart differences in biology of host-parasite interactions.

## INTRODUCTION

Parasites in the genus *Leishmania* cause a spectrum of disease ranging from cutaneous (CL) to visceralizing (VL) disease and are frequently used in many experimental animal models. Up to 1.6 million people each year are infected with one of the forms of leishmaniasis (1). This significant impact of disease resulted in an estimated 980,000 disability adjusted life years (DALYs) in 2016 (2). Further, these estimates are likely to significantly underestimate the burden caused by social and psychological stigmatization resulting from long term scarring (3).

The outcome of *Leishmania* infection has been understood to depend on proper T-helper (T_h_) cell polarization since the late 1980’s when it was shown that a T_h_1 response promotes a healing immune response whereas a Th2 response leads to progressive, non-healing disease (4, 5). Since then, animal models of leishmaniasis have continued to be used to characterize and understand many important nuances of the adaptive immune system. Despite these many advances in understanding immunologic concepts broadly, our understanding of the immune response to *Leishmania* spp. specifically is evolving to reflect greater nuances and complexities (6). Further, current treatment options for leishmaniasis remain prolonged, expensive, have variable efficacy, and significant side effects presenting an urgent need for novel therapeutics (7). In order to investigate these effectively, it is paramount to have optimal laboratory techniques for assessing disease progression in animals accurately, reproducibly, and under a variety of experimental conditions.

Methods to monitor disease progression during *Leishmania* infection in cutaneous animal models have traditionally relied on measuring footpad swelling and using a limiting dilution assay (LDA) to quantify parasite burden (8). While LDA remains a reliable and sensitive technique for quantifying parasite burden, these assays are labor intensive, can be technically challenging, and take several weeks to obtain final results. Recent advances employing genetic manipulation of parasites to express live reporter molecules such as mCherry or luciferase allow advanced monitoring of the parasite in real time and over a long-time course; however, they require an additional layer of genetic manipulation on the parasite and often require expensive equipment for visualization (9-11). More recently developed protocols utilizing amplification of nucleic acids have been optimized for use in clinical diagnosis. To meet the demands of a clinical diagnostic assay, such as high sensitivity and the ability to discriminate between *Leishmania* spp., highly variable and multi-copy genes are most commonly used.

However, characteristics for clinical assays for *Leishmania* detection are not necessarily ideal for use in experimental animal models. The ability to differentiate between different strains is not required when the infecting species is carefully controlled and delivered during experimental infection. Additionally, the use of multi-copy genes which are known to be present in regions of relative genomic plasticity, may change during the course of infection. This at best adds unpredictable variation to the assay and at worst confounds the change by inducing a systematic change across only certain experimental conditions (12). Finally, it is particularly important to select genes without an active role in disease, as any studies to further understand these or related pathways are subject to the risk of mutations or gene copy expansions rendering PCR at these sites unable to accurately compare across experimental conditions.

Recently, several efforts have been made to standardize lab protocols for detection and quantification of parasites from clinical isolates (13, 14). However, similar comparative studies that emphasize assay traits optimized for experimental models of infection are lacking. Here we report a real time PCR (RT-PCR) assay based on amplification of a single-copy, housekeeping *Leishmania* RNA binding protein (DRBD3) that is optimal for animal model studies. With touchdown cycling parameters, we were able to achieve a sensitivity of 100 fg per reaction which rivals most described PCR protocols for *Leishmania* quantification. Use of this assay in the future will facilitate studies elucidating mechanisms of immunity to *Leishmania* spp. and in monitoring efficacy of novel pharmaceutical interventions.

## Results

### *RT-PCR assay design for the* Leishmania *RNA binding protein, DRBD3*

To design an RT-PCR assay with optimal characteristics for quantification of *Leishmania* parasite burden in experimental models, we first sought to identify a single copy gene essential to the parasite, but less likely to be actively involved in evasion of host defense. For this we chose putative *Leishmania* RNA binding protein (DRBD3), present in a single-copy on chromosome 4. The gene is classified as the double RNA binding domain 3 protein based on homology with *DRBD3* in *Trypanosoma brucei.* Studies in *T. brucei* have identified a consensus binding sequence in the 3’UTR of mRNA transcripts and suggest that DRBD3 plays a role in modulating stress response (15, 16); however, there have been no functional studies of this protein in *Leishmania* spp. to date. The *L. major* entry for DRBD3 (LmjF.04.1170) on TriTrypDB indicates that the gene is constitutively expressed between parasite life-cycle stages, is not under immune pressure, and has minimal sequence variation between species (17). RNA-seq experiments in both *L. major* and *L. amazonensis* have demonstrated little to no change in expression of this gene between the promastigote and amastigote stage consistent with the characteristics of a constitutively expressed housekeeping gene (18-20). Additionally, there were no identified epitopes in the Immune Epitope Database (IEDB) corresponding to DRBD3 peptides, suggesting that it is not likely to be influenced by host immune pressure (21). To verify the homology of DRBD3 between *Leishmania* spp. we used the sequence from *L. major Sd.* as a reference, and used NCBI blast software to identify all related sequences. This blast search yielded 11 hits of protein coding genomic DNA within the genus *Leishmania*. Using ClustalOmega we then created a multisequence alignment, and in combination with NCBI Primer Blast tool designed primers over a highly-conserved region (Figure 1A). Both primers have complete conservation at 18 out of 20 nucleotides. This design allows using a single set of primers to PCR amplify a stable, housekeeping gene that is likely to be applicable across a broad range of *Leishmania* spp.

**Figure 1.**
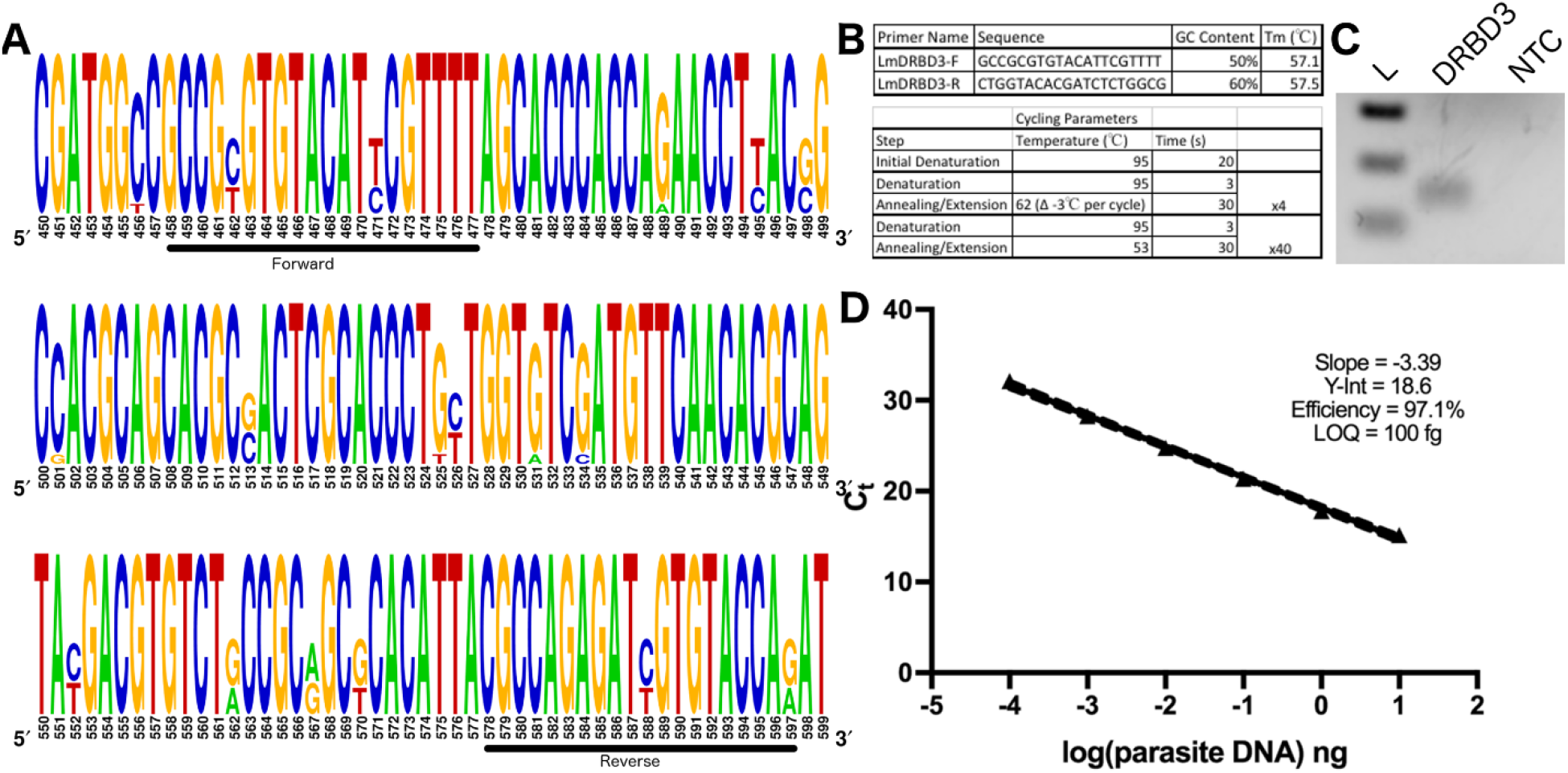
Primer design and optimization of DRBD3 based RT-PCR for parasite quantification. A) Multisequence alignment based on 11 homologous sequences to *L. major* Sd. DRBD3 found using NCBI Blast B) Primer sequences and cycling parameters used. C). DRBD3 primers amplify a 150bp product specifically. Product visualized with ethidium bromide staining of a 1% agarose gel run. D) Plot of Ct value vs log dilution of parasite burden to calculate primer efficiency.

The assay was then validated with DNA from cultured *L. major* promastigotes. With touchdown PCR cycling parameters, one distinct product is visible by ethidium bromide staining, which corresponds with a single product by melt curve analysis at 84.4°C (Figure 1B-C). Based on a standard curve extending across six orders of magnitude from 1×10^7^ fg – 100 fg of parasite DNA per reaction, the PCR had an average efficiency of 100.8% (96.98-104.7, 95%CI, n=6) (Figure 1D). The assay can detect DNA above this, but is no longer within the linear range. Therefore, we successfully developed a novel touch-down based PCR assay for a single-copy, housekeeping gene in *Leishmania* spp. that is able to efficiently detect as low as 100 fg of parasite DNA per reaction.

### Validation of DRBD3 RT-PCR in vivo

We then tested the assay’s ability to monitor parasite burden in the murine model of cutaneous leishmaniasis. Wildtype C57BL6 mice were inoculated subcutaneously with 2×10^6^ cultured promastigotes in the left hind footpad. At 1 day and 34 days post infection, DNA was isolated from the infected and uninfected footpads as well as the corresponding popliteal lymph nodes.

The assay was able to accurately monitor parasite burden over this course of infection. Reactions with DNA from uninfected footpads and non-draining lymph nodes, included as negative controls, did not amplify a single product at the corresponding melt temperature (84.7°C) in any sample (Figure 2A). At 1 day post infection, 7 of 7 mice had low parasite burdens in the footpad, but only 5 out of 7 infected mice had detectable parasites in the draining lymph node. This is consistent with an expected delay in time for migration of *Leishmania* parasites to the draining lymph nodes. In both the draining lymph nodes and infected footpads, a significant increase in parasite burden was observed between 1 day and 34 days post infection (Figure 2C-D). Therefore, we report that this assay successfully monitors *Leishmania* parasite burden in a mouse model of cutaneous leishmaniasis.

**Figure 2.**
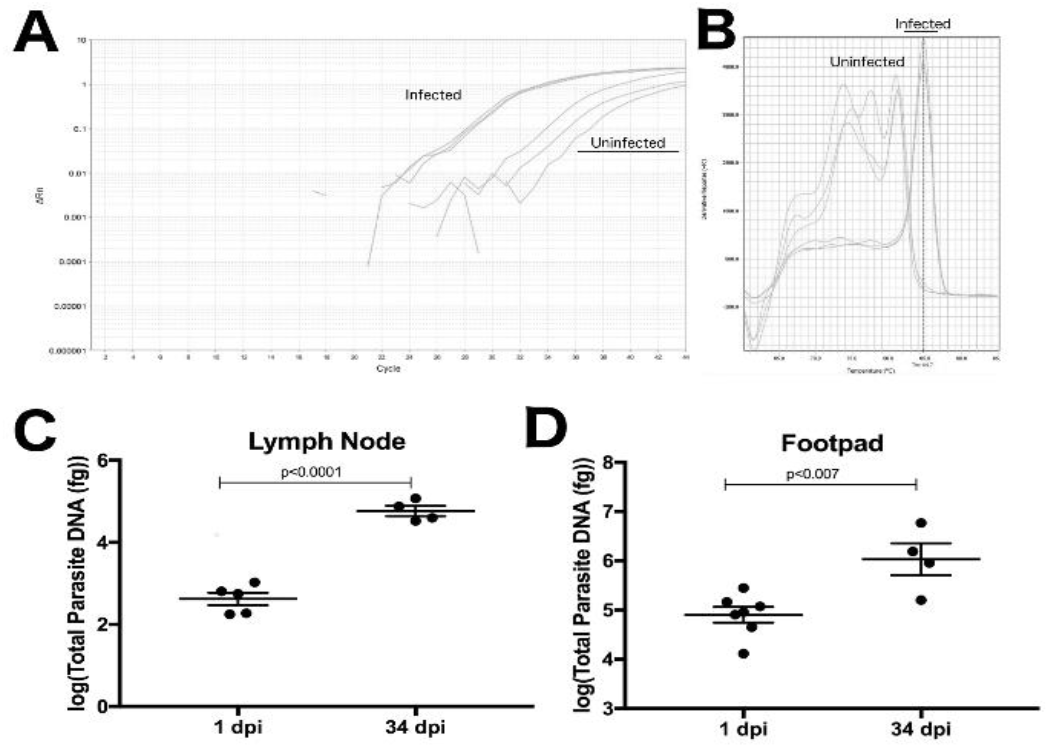
DRBD3 primers are able to assess parasite burden from infected mouse tissue. A-B) Representative amplification plots and melt curve analysis from the draining lymph nodes of mice at 34 days post infection. Parasites amplified from infected tissue had melting temperatures corresponding with the expected 84.7°C. Uninfected tissue produced nonspecific products with melt temps at lower temperatures. C) Quantification of total parasites in the draining popliteal lymph nodes at 1 and 34 days post infection. At 1 day post infection, 2 mice are not included because there was no detectable parasite burden at that time. D) Quantification of total parasites in the infected footpad at 1 and 34 days post infection. P-values calculated by parametric Students T-test.

### *Comparison of DRBD3 RT-PCR assay to other targets for quantification of* Leishmania spp. *infection in animal models*

We systematically compared the characteristics of the DRBD3 assay to other RT-PCR assays. A wide variety of assays for *Leishmania* detection by PCR with diverse targets and characteristics have been described; though, most have been optimized from the perspective of clinical diagnostics.

The most commonly used assays target multicopy genes. This produces advantages in regards to lower limits of detection and the potential to discriminate between species but come at the cost of uncertainty in gene stability at these plastic regions of the genome. 18s rRNA is present in up to 166 copies per parasite (22, 23), ITS-1 at 20-200 (still looking for original paper), minicircle kinetoplast DNA at up to 10,000 copies (24), and HSP70 at 5-7 (25). These assays report limits of detection as low as 10 fg of parasite DNA per reaction. However, multicopy regions are often under significant change. For example, kDNA is particularly unstable in terms of copy number with reports of it varying between species, strains, and even lifecycle stages (26, 27).

Single copy genes tend to be in more stable regions of the genome, but report higher limits of detection compared to multicopy gene assays and variable efficiencies between studies. Glucosephosphate isomerase (GPI), glucose 6 phosphate dehydrogenase (G6PD), Superoxide Dismutase 1 (SODB1) (28), Arginine Permease (AAP3) (29), and DNA polymerase alpha (30) are all single copy genes for which *Leishmania* quantification by PCR has been described. The assay for GPI reports a higher limit of detection of 5.6 pg compared to other assays with limits of detection around 10-100 fg per reaction (31). Reported PCR efficiencies for the G6PD assays range from 50.4% to 95.7%. SODB1 and AAP3 described RT-PCR assays have competitive limits of detection and efficiencies (28, 29); however, it has become increasingly apparent that both of these genes are important virulence factors. AAP3 is upregulated in response to arginine shortages in host macrophages, and SODB1 deficient *L. chagasi* parasites demonstrate impaired survival in host macrophages (32-35). This raises concern about the reliability of the assays during experimental manipulation; particularly in light of *Leishmania* spp. regulating gene expression through copy number variation (36).

Amplifying the gene target DNA polymerase alpha has similar characteristics to the DRBD3 assay. It is another example of single copy gene, which can be amplified at high efficiency with a limit of detection of 100 fg (37, 38). Therefore, for monitoring parasite burden accurately and precisely in animal models, the DRBD3 RT-PCR assay and the DNA polymerase alpha assay (38) fulfill the optimal characteristics.

**Table 1.**
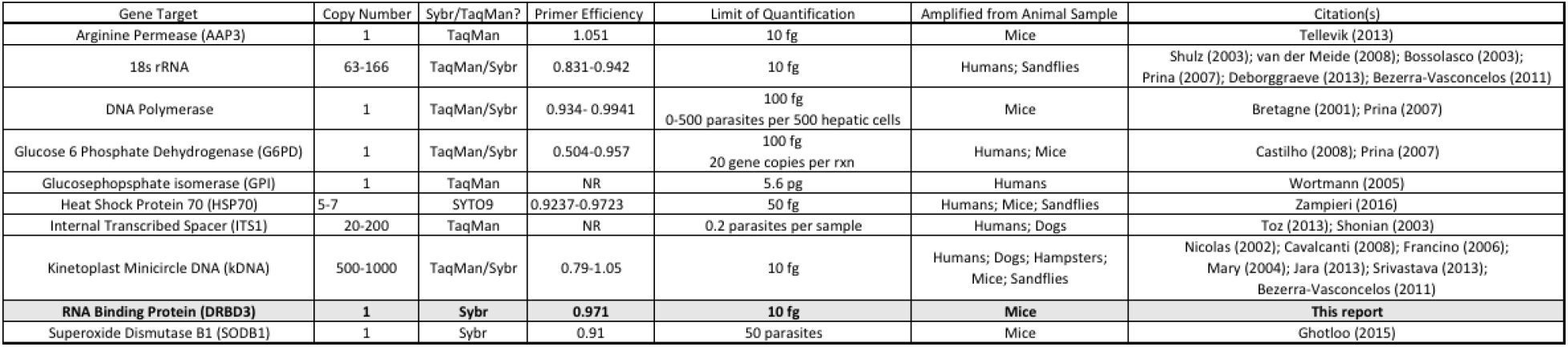
Comparison of described targets for RT-PCR based quantification of parasite burden in humans and animals.

## Discussion

Animal models of leishmaniasis have proven to be valuable in understanding basic immunological concepts, and will be critical in future drug development programs to control this neglected tropical disease. The ability to accurately monitor parasite survival and replication in these models is paramount to properly understanding the biology of infection and monitoring the effectiveness of novel therapeutic interventions. The development of RT-PCR assays for parasite quantification has been widespread for clinical applications; however, there is no standardized RT-PCR protocol that is optimized specifically for use in animal models. Here we developed a novel touchdown RT-PCR assay of the single copy housekeeping gene, DRBD3, which has a limit of detection that rivals existing protocols while not being subject to changes in sequence or copy number that multicopy genes or genes required for virulence would be subject to.

Accurate measurement of parasite burden by RT-PCR is important for distinguishing between host and parasite mediated pathology. Disease progression of Leishmania animal models is traditionally done by a combination of measuring footpad thickness or directly measuring parasites through limiting dilution assays. Measuring the inflammation that results from infection by swelling at the site of inoculation is a valuable way to monitor severity of disease, but it is unable to distinguish between damage caused by overgrowth of the parasite and damage caused by dysregulated host derived inflammation. Studies showing that parasite growth does not perfectly correlate with lesion size, demonstrate the importance of distinguishing between parasite and host mediated pathology (37, 39). This is particularly relevant in the context of leishmaniasis where it is well documented that distinct disease manifestations are caused by both extremes of this spectrum (40). Limiting dilution assays are useful for enumerating parasite burden, but are costly in terms of time and resources. Using a RT-PCR approach to amplify parasite DNA allows determination of parasite burden in as little as 2-4 hours instead of waiting approximately a week for parasites to grow in the LDA.

The single copy gene DRBD3 is highly conserved across *Leishmania* spp., and the primers used in this assay bind to a region of DNA over which 90% of the residues are completely conserved. However, it should be noted that the divergence of the 2 nucleotide positions occurs at the division between the *Leishmania leishmania* and *Leishmania viannia* subgenera. The primers reported here are 100% conserved within the *leishmania* subgenus. Within the *viannia* subgenus, the sequences are also 100% conserved, and are only altered at two sites between the two genera for each primer. We postulate that the primers are also likely to work with high efficiency for parasites in the *viannia* subgenus, or the assay would be easily adaptable to the *viannia* subgenus with a second set of specific primers. The DRBD3 RT-PCR assay for *Leishmania* quantification is therefore also likely to be a robust assay for labs in that validation of the assay one time will allow for use in a wide range of studies modeling parasites with distinct disease phenotypes and from diverse geographic and evolutionary backgrounds.

Careful consideration of protocols used to quantify parasite burden in experimental models of leishmaniasis is essential to fully understand host-parasite interactions and for assessing the efficacy of novel therapeutic interventions. The DRBD3 assay described here will allow for consistent detection of parasites in an unbiased manner with a low limit of detection, facilitating discoveries in basic science and improvements in leishmaniasis treatment.

## Materials and Methods

### Multisequence alignment and sequence Logo

Traits of the DRBD3 gene were analyzed in TriTrypDB using the L. major Fd reference sequence (LmjF.04.1170) (17). The reference sequence used in NCBI Blast tool to identify homologous sequences was based on the *L. major* SD75.1 sequence in order to correspond with the parasite DNA used to validate the assay in this study. The 11 identified homologous genomic DNA sequences were downloaded and aligned using ClustalOmega (41). The WebLogo tool was used to generate a sequence logo based on this alignment (http://weblogo.berkeley.edu/) (42).

### Primer design

Primers were designed using Primer-BLAST software (43). The gene for DRBD3 in the *Leishmania major* (MHOM/SN/74/Seidman) was input. Five pairs of primers with similar melt temperatures were initially tested. After amplification by conventional PCR, 3 of these primer pairs resulted in non-specific amplification as detected by ethidium bromide detection in a 1% agarose gel. Based on this initial screen, the primers reported in Figure 1B were then used to amplify wildtype *L. major* (MHOM/SN/74/Seidman) promastigote DNA as described below.

### RT-PCR protocol for parasite quantification

After DNA isolation from each mouse tissue, 100ng of DNA was used per reaction on a plate with a standard curve of cultured promastigote DNA ranging from 1X10^7^ fg – 100 fg per reaction. PCR reactions were set up in a final volume of 10μl by adding 5μl of ITaq universal SYBR Green supermix (BioRad, Cat# 172-5121), 0.5μl of each primer (1μM concentration), and 100ng genomic DNA per sample. Cycling parameters are described in Figure 1B. All PCR reactions were performed in triplicate. Reactions were excluded from the final analysis if by melt temperature analysis there was any peak outside of the expected 84.7°C product. Primer efficiency for each reaction was calculated using the formula 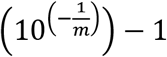, where *m* is the slope from a plot of the Ct vs log(parasite DNA) with serial 10 fold dilutions ranging from 1×10^7^fg to 100fg total DNA per reaction.

Total parasite DNA was calculated by first using the Ct for each reaction to interpolate the fg of parasite DNA per 100ng of total DNA relative to the standard curve included on each 96-well plate. DNA (fg per reaction) was then multiplied by the factor used to dilute each sample to 100ng per PCR reaction in order to get total parasite DNA (fg) per tissue harvested.

### Parasites and culture conditions

*Leishmania major* (MHOM/SN/74/Seidman) were obtained from BEI (BEI reference# 48819). Parasites were maintained in M199 media supplemented with 10% heat inactivated FBS and 0.2% hemin. Cultures were maintained by inoculating a new 10mL culture with 200ul of previous culture every 5 days. Infections in mice were performed with parasites that had been passaged less than 3 times *in vitro.*

### Mouse infections

Wildtype C57BL6 mice from Jackson were maintained in the Duke Laboratory Animal Resource (DLAR). All studies are approved under Duke University IACUC protocol A200-15-7. *L. major* (MHOM/SN/74/Seidman) parasites were prepared by washing 5 day old culture of promastigotes with Hanks Buffered Salt Solution (HBSS), counting by hemocytometer, and resuspending at 2 × 10^7^ parasties per 50μl of HBSS. A 27G 1/2 mL with permanently attached needle was used to inoculate the left hind footpad with 50μl of promastigote suspension. Mice were monitored at least twice weekly to track lesion development.

DNA was obtained using the Qiagen DNeasy Blood and Tissue Kit (Cat # 69504). In brief, tissue from the infected footpads and draining popliteal lymph node of each mouse was harvested. The contralateral footpad and lymph node was taken from each mouse to monitor for contamination. Tissue was placed into a clean 1.5mL microcentrifuge tube. To 180μl of buffer ATL and 20μl of proteinase K was added to each tube prior to homogenizing the tissue with a bead beater and incubating samples at 37°C overnight before proceeding as indicated by manufacturer instructions. DNA quantity and quality was assessed using the Take3 application on a Synergy H1 BioTek plate reader prior to use in the RT-PCR assay described.

## References

1. Alvar J, Velez ID, Bern C, Herrero M, Desjeux P, Cano J, Jannin J, den Boer M, Team WHOLC. 2012. Leishmaniasis worldwide and global estimates of its incidence. PLoS One 7:e35671.

2. DALYs GBD, Collaborators H. 2017. Global, regional, and national disability-adjusted life-years (DALYs) for 333 diseases and injuries and healthy life expectancy (HALE) for 195 countries and territories, 1990-2016: a systematic analysis for the Global Burden of Disease Study 2016. Lancet 390:1260–1344.

3. Bailey F, Mondragon-Shem K, Hotez P, Ruiz-Postigo JA, Al-Salem W, Acosta-Serrano A, Molyneux DH. 2017. A new perspective on cutaneous leishmaniasis-Implications for global prevalence and burden of disease estimates. PLoS Negl Trop Dis 11:e0005739.

4. Heinzel FP, Sadick MD, Holaday BJ, Coffman RL, Locksley RM. 1989. Reciprocal expression of interferon gamma or interleukin 4 during the resolution or progression of murine leishmaniasis. Evidence for expansion of distinct helper T cell subsets. J Exp Med 169:59–72.

5. Scott P, Natovitz P, Coffman RL, Pearce E, Sher A. 1988. CD4+ T cell subsets in experimental cutaneous leishmaniasis. Mem Inst Oswaldo Cruz 83 Suppl 1:256–259.

6. Scott P, Novais FO. 2016. Cutaneous leishmaniasis: immune responses in protection and pathogenesis. Nat Rev Immunol doi:10.1038/nri.2016.72.

7. Ponte-Sucre A, Gamarro F, Dujardin JC, Barrett MP, Lopez-Velez R, Garcia-Hernandez R, Pountain AW, Mwenechanya R, Papadopoulou B. 2017. Drug resistance and treatment failure in leishmaniasis: A 21st century challenge. PLoS Negl Trop Dis 11:e0006052.

8. Sacks DL, Melby PC. 2001. Animal models for the analysis of immune responses to leishmaniasis. Curr Protoc Immunol Chapter 19:Unit 19 12.

9. Michel G, Ferrua B, Lang T, Maddugoda MP, Munro P, Pomares C, Lemichez E, Marty P. 2011. Luciferase-expressing Leishmania infantum allows the monitoring of amastigote population size, in vivo, ex vivo and in vitro. PLoS Negl Trop Dis 5:e1323.

10. Calvo-Alvarez E, Guerrero NA, Alvarez-Velilla R, Prada CF, Requena JM, Punzon C, Llamas MA, Arevalo FJ, Rivas L, Fresno M, Perez-Pertejo Y, Balana-Fouce R, Reguera RM. 2012. Appraisal of a Leishmania major strain stably expressing mCherry fluorescent protein for both in vitro and in vivo studies of potential drugs and vaccine against cutaneous leishmaniasis. PLoS Negl Trop Dis 6:e1927.

11. Roy G, Dumas C, Sereno D, Wu Y, Singh AK, Tremblay MJ, Ouellette M, Olivier M, Papadopoulou B. 2000. Episomal and stable expression of the luciferase reporter gene for quantifying Leishmania spp. infections in macrophages and in animal models. Mol Biochem Parasitol 110:195–206.

12. Laffitte MN, Leprohon P, Papadopoulou B, Ouellette M. 2016. Plasticity of the Leishmania genome leading to gene copy number variations and drug resistance. F1000Res 5:2350.

13. Cruz I, Millet A, Carrillo E, Chenik M, Salotra P, Verma S, Veland N, Jara M, Adaui V, Castrillon C, Arevalo J, Moreno J, Canavate C. 2013. An approach for interlaboratory comparison of conventional and real-time PCR assays for diagnosis of human leishmaniasis. Exp Parasitol 134:281–289.

14. Leon CM, Munoz M, Hernandez C, Ayala MS, Florez C, Teheran A, Cubides JR, Ramirez JD. 2017. Analytical Performance of Four Polymerase Chain Reaction (PCR) and Real Time PCR (qPCR) Assays for the Detection of Six Leishmania Species DNA in Colombia. Front Microbiol 8:1907.

15. Fernandez-Moya SM, Garcia-Perez A, Kramer S, Carrington M, Estevez AM. 2012. Alterations in DRBD3 ribonucleoprotein complexes in response to stress in Trypanosoma brucei. PLoS One 7:e48870.

16. Das A, Bellofatto V, Rosenfeld J, Carrington M, Romero-Zaliz R, del Val C, Estevez AM. 2015. High throughput sequencing analysis of Trypanosoma brucei DRBD3/PTB1-bound mRNAs. Mol Biochem Parasitol 199:1–4.

17. Aslett M, Aurrecoechea C, Berriman M, Brestelli J, Brunk BP, Carrington M, Depledge DP, Fischer S, Gajria B, Gao X, Gardner MJ, Gingle A, Grant G, Harb OS, Heiges M, Hertz-Fowler C, Houston R, Innamorato F, Iodice J, Kissinger JC, Kraemer E, Li W, Logan FJ, Miller JA, Mitra S, Myler PJ, Nayak V, Pennington C, Phan I, Pinney DF, Ramasamy G, Rogers MB, Roos DS, Ross C, Sivam D, Smith DF, Srinivasamoorthy G, Stoeckert CJ, Jr., Subramanian S, Thibodeau R, Tivey A, Treatman C, Velarde G, Wang H. 2010. TriTrypDB: a functional genomic resource for the Trypanosomatidae. Nucleic Acids Res 38:D457–462.

18. Aoki JI, Muxel SM, Zampieri RA, Laranjeira-Silva MF, Muller KE, Nerland AH, Floeter-Winter LM. 2017. RNA-seq transcriptional profiling of Leishmania amazonensis reveals an arginase-dependent gene expression regulation. PLoS Negl Trop Dis 11:e0006026.

19. Akopyants NS, Matlib RS, Bukanova EN, Smeds MR, Brownstein BH, Stormo GD, Beverley SM. 2004. Expression profiling using random genomic DNA microarrays identifies differentially expressed genes associated with three major developmental stages of the protozoan parasite Leishmania major. Mol Biochem Parasitol 136:71–86.

20. Leifso K, Cohen-Freue G, Dogra N, Murray A, McMaster WR. 2007. Genomic and proteomic expression analysis of Leishmania promastigote and amastigote life stages: the Leishmania genome is constitutively expressed. Mol Biochem Parasitol 152:35–46.

21. Vita R, Zarebski L, Greenbaum JA, Emami H, Hoof I, Salimi N, Damle R, Sette A, Peters B. 2010. The immune epitope database 2.0. Nucleic Acids Res 38:D854–862.

22. Ivens AC, Peacock CS, Worthey EA, Murphy L, Aggarwal G, Berriman M, Sisk E, Rajandream MA, Adlem E, Aert R, Anupama A, Apostolou Z, Attipoe P, Bason N, Bauser C, Beck A, Beverley SM, Bianchettin G, Borzym K, Bothe G, Bruschi CV, Collins M, Cadag E, Ciarloni L, Clayton C, Coulson RM, Cronin A, Cruz AK, Davies RM, De Gaudenzi J, Dobson DE, Duesterhoeft A, Fazelina G, Fosker N, Frasch AC, Fraser A, Fuchs M, Gabel C, Goble A, Goffeau A, Harris D, Hertz-Fowler C, Hilbert H, Horn D, Huang Y, Klages S, Knights A, Kube M, Larke N, Litvin L, et al. 2005. The genome of the kinetoplastid parasite, Leishmania major. Science 309:436–442.

23. Torres-Machorro AL, Hernandez R, Cevallos AM, Lopez-Villasenor I. 2010. Ribosomal RNA genes in eukaryotic microorganisms: witnesses of phylogeny? FEMS Microbiol Rev 34:59–86.

24. Yurchenko VY, Merzlyak EM, Kolesnikov AA, Martinkina LP, Vengerov YY. 1999. Structure of Leishmania minicircle kinetoplast DNA classes. J Clin Microbiol 37:1656–1657.

25. Folgueira C, Canavate C, Chicharro C, Requena JM. 2007. Genomic organization and expression of the HSP70 locus in New and Old World Leishmania species. Parasitology 134:369–377.

26. Weirather JL, Jeronimo SM, Gautam S, Sundar S, Kang M, Kurtz MA, Haque R, Schriefer A, Talhari S, Carvalho EM, Donelson JE, Wilson ME. 2011. Serial quantitative PCR assay for detection, species discrimination, and quantification of Leishmania spp. in human samples. J Clin Microbiol 49:3892–3904.

27. Mary C, Faraut F, Lascombe L, Dumon H. 2004. Quantification of Leishmania infantum DNA by a real-time PCR assay with high sensitivity. J Clin Microbiol 42:5249–5255.

28. Ghotloo S, Haji Mollahoseini M, Najafi A, Yeganeh F. 2015. Comparison of Parasite Burden Using Real-Time Polymerase Chain Reaction Assay and Limiting Dilution Assay in Leishmania major Infected Mouse. Iran J Parasitol 10:571–576.

29. Tellevik MG, Muller KE, Lokken KR, Nerland AH. 2014. Detection of a broad range of Leishmania species and determination of parasite load of infected mouse by real-time PCR targeting the arginine permease gene AAP3. Acta Trop 137:99–104.

30. Croan DG, Morrison DA, Ellis JT. 1997. Evolution of the genus Leishmania revealed by comparison of DNA and RNA polymerase gene sequences. Mol Biochem Parasitol 89:149–159.

31. Wortmann G, Hochberg L, Houng HH, Sweeney C, Zapor M, Aronson N, Weina P, Ockenhouse CF. 2005. Rapid identification of Leishmania complexes by a real-time PCR assay. Am J Trop Med Hyg 73:999–1004.

32. Castilho-Martins EA, Laranjeira da Silva MF, dos Santos MG, Muxel SM, Floeter-Winter LM. 2011. Axenic Leishmania amazonensis promastigotes sense both the external and internal arginine pool distinctly regulating the two transporter-coding genes. PLoS One 6:e27818.

33. Muxel SM, Laranjeira-Silva MF, Zampieri RA, Floeter-Winter LM. 2017. Leishmania (Leishmania) amazonensis induces macrophage miR-294 and miR-721 expression and modulates infection by targeting NOS2 and L-arginine metabolism. Sci Rep 7:44141.

34. Goldman-Pinkovich A, Balno C, Strasser R, Zeituni-Molad M, Bendelak K, Rentsch D, Ephros M, Wiese M, Jardim A, Myler PJ, Zilberstein D. 2016. An Arginine Deprivation Response Pathway Is Induced in Leishmania during Macrophage Invasion. PLoS Pathog 12:e1005494.

35. Plewes KA, Barr SD, Gedamu L. 2003. Iron superoxide dismutases targeted to the glycosomes of Leishmania chagasi are important for survival. Infect Immun 71:5910–5920.

36. Iantorno SA, Durrant C, Khan A, Sanders MJ, Beverley SM, Warren WC, Berriman M, Sacks DL, Cotton JA, Grigg ME. 2017. Gene Expression in Leishmania Is Regulated Predominantly by Gene Dosage. MBio 8.

37. Bretagne S, Durand R, Olivi M, Garin JF, Sulahian A, Rivollet D, Vidaud M, Deniau M. 2001. Real-time PCR as a new tool for quantifying Leishmania infantum in liver in infected mice. Clin Diagn Lab Immunol 8:828–831.

38. Prina E, Roux E, Mattei D, Milon G. 2007. Leishmania DNA is rapidly degraded 410 following parasite death: an analysis by microscopy and real-time PCR. Microbes Infect 9:1307–1315.

39. Hill JO, North RJ, Collins FM. 1983. Advantages of measuring changes in the number of viable parasites in murine models of experimental cutaneous leishmaniasis. Infect Immun 39:1087–1094.

40. Scott P, Novais FO. 2016. Cutaneous leishmaniasis: immune responses in protection and pathogenesis. Nat Rev Immunol 16:581–592.

41. McWilliam H, Li WZ, Uludag M, Squizzato S, Park YM, Buso N, Cowley AP, Lopez R. 2013. Analysis Tool Web Services from the EMBL-EBI. Nucleic Acids Research 41:W597–W600.

42. Crooks GE, Hon G, Chandonia JM, Brenner SE. 2004. WebLogo: A sequence logo generator. Genome Research 14:1188–1190.

43. Ye J, Coulouris G, Zaretskaya I, Cutcutache I, Rozen S, Madden TL. 2012. Primer-BLAST: a tool to design target-specific primers for polymerase chain reaction. BMC Bioinformatics 13:134.

